# Relative fire-proneness of land cover types in the Brazilian Atlantic Forest

**DOI:** 10.1101/2024.07.21.604452

**Authors:** Bruno FCB Adorno, Augusto AJ Piratelli, Erica Hasui, Milton C Ribeiro, Pedro G. Vaz

## Abstract

Fires are increasingly affecting tropical biomes, where landscape-fire interactions remain understudied. We investigate the fire-proneness—the likelihood of a land use or land cover (LULC) type burning more or less than expected based on availability—in the Brazilian Atlantic Forest (AF). This biodiversity hotspot is increasingly affected by fires due to human activities and climate change. Using a selection ratio-based approach, we analyzed fire-LULC interactions in 40,128 fires over a 35-year period (1987-2022) across various ecoregions in the AF. Our findings revealed that secondary forests, forest areas that have regrown after major disturbances, burned 61% more than expected by chance, whereas old-growth forests, native forests that have developed over very long periods, burned 57% less than expected, highlighting a nearly inverse relationship in their fire-proneness. Interestingly, our data indicate that pastures in the AF are less prone to fire than expected, despite being considered among the land uses that burn the most in Brazil. Other LULCs showed variable fire-proneness, with some differences between ecoregions. Over time, the fire-proneness of secondary forests decreased, likely due to forest aging and changes in land management practices. We emphasize the necessity for tailored fire management strategies that address the unique vulnerabilities of secondary forests, particularly in the context of ongoing restoration efforts aimed at increasing native forests. Effective measures, including the implementation of ‘fire-smart management’ practices and enhancing the perceived value of secondary forests among local communities, are crucial for mitigating fire risks. Integrating these strategies with incentive-based approaches can bolster fire prevention, ensuring the long-term success of restoration programs. Our study provides a framework for understanding fire-landscape dynamics in tropical forests and offers actionable insights for practitioners working to safeguard these biomes from the escalating threat of wildfires.

**GRAPHICAL ABSTRACT:** 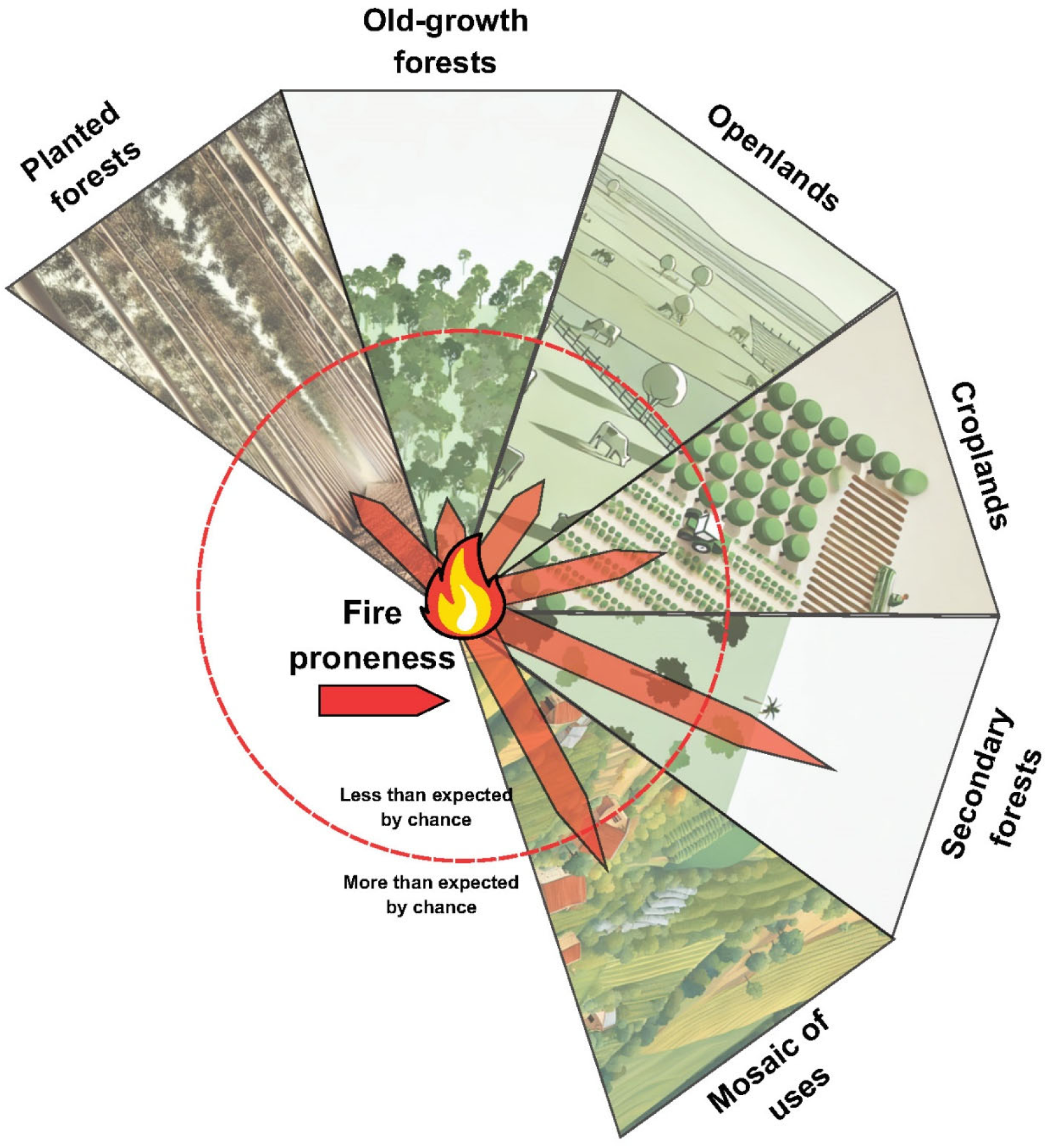

**Highlights:**

- Evaluation of land cover fire-proneness in a tropical biome using a selection ratio.
- Secondary forests burned 61% more; old-growth forests 57% less than expected by chance.
- Pastures in the Atlantic Forest seem less fire-prone than expected, despite frequent burns.
- Fire-proneness of secondary forests has generally been decreasing over time.
- Tailored fire management is essential to protect secondary forests in restoration efforts.

## 1. INTRODUCTION

Tropical forests are fire-sensitive biomes, with species lacking evolutionary adaptations to frequent fires and other aspects of fire regimes (Hoffmann et al., 2012; Pivello et al., 2021). Historically, these biomes experienced fires every few hundred years (Bush et al., 2008), but climate change and human activities have altered the frequency and nature of these events (Cochrane, 2003; Fonseca et al., 2019; Le Page et al., 2017). Fires now occur every decade or even annually, often escalating in size and severity (Cochrane, 2003; Flannigan et al., 2009; Uhl & Kauffman, 1990). Fires in tropical forests typically occur as low-severity surface fires, which consume the dry litter layer and result in the death of small trees and seedlings (Barlow et al., 2002; Staver et al., 2020; Uhl & Kauffman, 1990). These surface fires can escalate to high-severity fires, killing most large trees with thin bark. The resulting canopy openings increase sunlight penetration, which dries out the forest and creates conditions favorable to subsequent fires (Sansevero et al., 2020; Pivello et al., 2021). Hence, fires are associated with biodiversity changes (Kelly et al., 2017; Kelly et al, 2020), soil degradation (Nadporozhskaya et al., 2018), erosion (Girona-García et al., 2024), impacts on freshwater ecosystems (Vaz et al., 2014, 2021), diminished water retention (Sansevero et al., 2017; Schmerbeck & Fiener, 2015), and landscape dynamics alterations (Silva et al., 2011; Moreira et al., 2011).

Fires are particularly worrying in the Atlantic Forest (AF) biome, one of the world’s most important biodiversity hotspots, which has high endemism (Myers et al., 2000). Originally, the AF covered 160 million hectares across coastal and inland portions of Brazil, Argentina, and Paraguay (Muylaert et al., 2018). This biome is the most degraded and deforested Brazilian biome, having suffered from intense land use and land cover (LULC) changes in recent centuries (Rezende et al., 2018). Most old-growth forests—native forests that have developed over long periods without significant disturbance—were cleared to make way for human activities. Today, the AF retains only about 36% of its extent of natural vegetation. Most of this vegetation is highly fragmented, with 97% of forest patches being under 50 ha, isolated, and impacted by various human-induced disturbances (Ribeiro et al., 2009a; Vancine et al., 2024). Despite these challenges, recent decades have seen improvements in vegetation cover due to conservation efforts led by the government and environmental organizations, aimed at reducing deforestation rates (Piffer et al., 2022a). Also, the abandonment of agriculture and the depopulation of rural areas contributed to the resurgence of natural vegetation, particularly in the form of secondary forests—areas of forest that have regrown after major disturbances such as deforestation and fires, whose ecological value and recovery potential depend on the time since disturbance and the surrounding landscape context (Rezende et al., 2018; Souza et al., 2020; Vancine et al., 2024). Changes in Brazilian legislation, such as the Atlantic Forest Law in 2006 and the Brazilian Forest Code in 2012, are also expected to influence the fire-landscape dynamic over the years. However, despite strict regulations, fire is increasingly shaping the AF landscape (dos Santos et al., 2019), namely by resetting the gradual transitions from secondary forests to vegetation resembling old-growth forests. Fires disrupt regeneration processes and can shift the biome towards a savanna-like state (dos Santos et al., 2019; Sansevero et al., 2020).

Fires do not occur randomly across the LULC types of a given landscape; instead, their occurrence and spread are determined by a complex interplay of human actions and natural phenomena (Brando et al., 2016; Fonseca et al., 2019; Le Page et al., 2017; Pivello, 2011). In tropical forests, most fires are caused by humans as a result of using fire to clear or expand agricultural areas (da Silva Junior et al., 2020; Gois et al., 2020; Oliveira-Júnior et al., 2020). Fire spread through the landscape is influenced by wind, topography, vegetation structure, moisture content of the vegetation and soil, amount and composition of surface fuel load (Carmo et al., 2011; Moreira et al., 2009; Oliveira et al., 2014; Silva et al., 2009), and factors impacting firefighting efforts (de Assis Barros et al., 2022; Gutiérrez-Velez et al., 2014; Pivello et al., 2021). These factors interact with different LULC types, making some LULC types more susceptible to fire spread than others (Abreu et al., 2022; Fonseca et al., 2019; Gutiérrez-Vélez et al., 2014). We refer to this susceptibility as fire-proneness, reflecting the likelihood of a LULC type burning more or less frequently than expected based on its availability in the landscape. LULCs that are more fire-prone are likely to be disproportionately affected, beyond what would be expected by chance alone (Moreira et al., 2001).

In Brazilian tropical forests, fires disproportionately affect LULC types, with pastures and croplands being the most likely to burn (Abreu et al., 2022; Cano-Crespo et al., 2015; de Assis Barros et al., 2022; de Santana et al., 2020; Freitas et al., 2020; Gutiérrez-Vélez et al., 2014; Herrmann et al., 2023; Uriarte et al., 2016). Conversely, a negative correlation is expected between fire occurrence and spread and the cover of old-growth forests (de Assis Barros et al., 2022; Fonseca et al., 2019; Singh & Huang, 2022). This low fire-proneness among old-growth forests can be attributed to their remarkable retention of transpired moisture, which effectively reduces susceptibility to fire (Cochrane et al., 1999; Uhl et al., 1988). Unfortunately, ongoing transitions from old-growth forests to other types of LULC can often result in increased fire-proneness (de Assis Barros et al., 2022; Eva & Lambin, 2000; Fonseca et al., 2019; Singh & Huang, 2022). Particularly, changes from old-growth forests to secondary forests likely increase fire-proneness, as secondary forests are more vulnerable due to their differing species composition, lower moisture content, and elevated radiation and temperature conditions (Fonseca et al., 2019; Gutiérrez-Vélez et al., 2014). Secondary forests in the AF typically emerge in degraded, abandoned pasturelands unsuitable for agriculture and livestock, and may also regenerate on the edges of small forest fragments, thus increasing the risk of fire (Guedes et al., 2020). As secondary forests mature and develop characteristics typical of old-growth forests, their fire-proneness is expected to decrease (Lebrija-Trejos et al., 2011; Ray et al., 2010).

Despite the extensive research on landscape-wildfire dynamics, critical gaps remain in our understanding of how fire-proneness varies among different LULC within Tropical Forests. In Brazil, the bulk of fire research has been conducted in the Amazon Forest, leading to a disproportionate focus that overlooks the distinct dynamics within the AF (dos Santos et al., 2019; Sansevero et al., 2020). Furthermore, no previous studies have comprehensively assessed the fire-proneness of different LULCs in Brazilian Tropical forests (see Carmo et al., 2011; Moreira et al., 2009; Oliveira et al., 2014). Instead, research has typically focused on examining the distribution of burned areas and the number of fire foci (Abreu et al., 2022; de Assis Barros et al., 2022; Gutiérrez-Vélez et al., 2014). Moreover, existing studies often overlook the heterogeneity of ecoregions in the AF, their cultural and socio-economic differences, as well as the interannual variability and drivers of fire activity, which likely impact fire dynamics. The ecoregions, including evergreen (ombrophilous) and seasonal forests, as well as transitional ecotone areas, are shaped by distinct temperature and rainfall patterns (Joly et al., 2014). Their distribution and fire proneness are also influenced by the AF’s extensive longitudinal, latitudinal, and altitudinal variations (Marques et al., 2021). A lack of long-term studies further limits understanding of fire-LULC dynamics over time.

In this study, we used a selection ratio-based approach (e.g., Moreira et al., 2009) to analyze relative fire-LULC interactions across the ecoregions of the AF, examining data at five-year intervals over a 35-year period. By applying this established approach to tropical biomes, our study offers a novel assessment of fire-landscape relationships in this distinct region. We tested the following hypotheses. 1) Old-growth forests have low fire-proneness, irrespective of ecoregion or year analyzed. 2) Secondary forests exhibit high fire-proneness in all ecoregions compared to old-growth forests. 3) Secondary forests exhibit a decreasing trend in fire-proneness over the years as the average age of their patches increases, making them progressively more resemblant of old-growth forests, including more closed canopies and greater moisture at ground level. 4) Croplands, pastures, and other human-impacted LULCs exhibit variable fire-proneness dependent on ecoregion and year.

## 2. METHODS

We employed geospatial analyses to examine relative fire-proneness in different LULC types across the Brazilian AF ecoregions. Our workflow involved defining the ecoregions, obtaining and processing LULC and fire data, and estimating selection ratios to assess fire-proneness in each region.

### 2.1 Ecoregions

To assess how ecoregions affect fire-LULC interactions in the Brazilian AF, we analyzed the five regions that cover over 90% of the biome and record the most fires (Fig. 1). These ecoregions are defined by floristic compositions of typical genera and characteristic biological forms that recur within the same climate, potentially occurring on terrains with varied lithology but well-defined relief (IBGE, 2012). We utilized vector files from the Database of Environmental Information (IBGE; https://bdiaweb.ibge.gov.br), which employs the Brazilian Vegetation Classification to map and hierarchically organize the land cover across regions. The mapping methods are detailed in the IBGE’s Technical Manual of Brazilian Vegetation (IBGE, 2012). The ecoregions are described as follows:

1) Semideciduous seasonal forest (SSF): Covering 37% of the biome, this largest ecoregion in the AF is marked by two distinct seasons—rainy summers and dry winters. During the dry season, up to 50% of tree species undergo leaf shedding.
2) Decidual seasonal forest (DSF): Occupying 6% of the AF, these forests prevail at higher altitudes where temperatures are lower. Like the SSF, they experience two distinct seasons—rainy summers and dry winters. Over 50% of their tree species shed leaves during the dry season.
3) Dense ombrophilous forest (DOF): This region occupies 17% of the AF, forming a verdant belt along the Brazilian coast. Characterized by the absence of a dry season, it experiences consistently high precipitation year-round and maintains temperatures averaging above 25°C.
4) Mixed ombrophilous forest (MOF): Commonly known as ‘Araucaria forests’, these forests are notable for consistent high precipitation and a diverse mix of evergreen and deciduous species. Covering 15% of the AF, they are distinguished by a high prevalence of gymnosperm species, notably *Araucaria angustifolia*.
5) Contact areas (CA): Covering 15% of the AF, these undifferentiated plant communities represent transitional ecotone areas, which occur at the boundaries or within the interiors of different ecoregions. In the AF, these ecotones can be found between forests and open physiognomies, such as those typical of the Cerrado, as well as between different forest types, like evergreen and seasonal forests. These areas are often characterized by a mix of vegetation types with varying degrees of canopy closure, reflecting the transitional nature of the environmental gradients they occupy.

**Figure 1.**
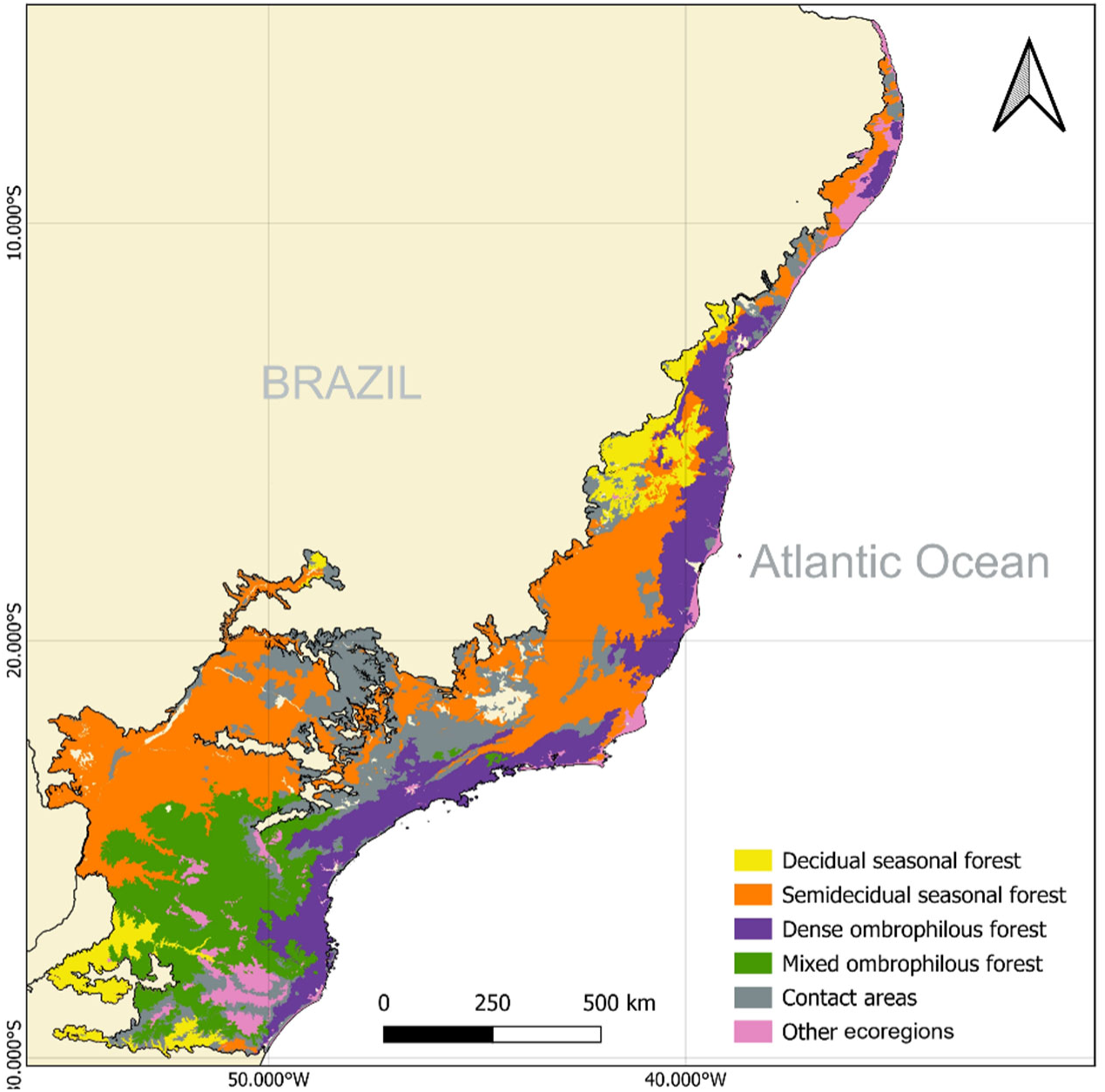
Study area. Ecoregions within the Brazilian Atlantic Forest.

### 2.2 Land cover maps

To obtain maps of LULC types in the ecoregions of the AF, we used maps from the MapBiomas platform - Collection 8.0 (available at https://brasil.mapbiomas.org). The MapBiomas Project is a collaborative Brazilian initiative that maps LULCs across the entire territory of Brazil. It uses satellite images, specifically from the Landsat program with a 30-m resolution, along with machine learning tools to create an annual time series of maps from 1985 to the present (Alencar et al., 2022). Transitions between classes are defined based on detectable changes in LULC, such as shifts from anthropogenic to forest classes, identified through algorithms that analyze satellite imagery. The platform also captures reversals, such as deforestation events, and secondary vegetation, where secondary vegetation is converted back to anthropogenic use. Additionally, the platform provides information on the age of secondary forests (Silva Junior et al., 2020).

We selected eight LULC maps, one for each five-year interval, spanning from 1986 to 2021 (Vancine et al., 2024). This interval helps minimize temporal autocorrelation, enhancing the independence of subsequent assessments of fire-LULC interactions. This same interval was previously used to assess vegetation dynamics in the AF (Vancine et al., 2024). Next, we simplified the 30 LULC classes from MapBiomas Collection 8.0 into seven categories (Table 1), plus a ‘Noncombustible areas’ type, which was omitted from further analysis. This reclassification was based on vegetation structural similarities and potential for comparable fire behavior, particularly in terms of fuel structure and composition (Nunes et al., 2005).

**Table 1.** Characterization of land use and land cover types.

| Land cover type | Definition | Land cover type in MapBiomass |
| --- | --- | --- |
| Old-growth forests | Areas with native forest cover since the beginning of the historical MapBiomass series. | Forest Formation; Floodable Forest |
| Secondary forests | Areas that have transitioned from anthropogenic use to native forest vegetation. The transition period (vegetation age) can vary from 1 to 35 years. | The captions for secondary forest areas provide information about the age of the vegetation. In this context, we grouped all age ranges into a single category corresponding to secondary forests. |
| Openlands | Areas characterized by ground and shrub vegetation. | Savanna Formation; Herbaceous Sandbank Vegetation; Grassland; Pasture; Other Non-forest Formations |
| Temporary crops | Areas that are both sown and harvested within the same agricultural year, sometimes more than once. | Soybean; Sugar cane; Rice; Cotton; Other Temporary Crops |
| Perennial crops | Areas with plant species that are cultivated and live for more than two years without needing to be replanted annually | Coffee; Citrus; Palm Oil; Other Perennial Crops |
| Planted forests | Areas dedicated to silviculture, primarily consisting of *Eucalyptus* sp. and *Pinus* sp. | Forest Plantation |
| Mosaic of uses | Mix or combination of different land uses, creating a diverse and complex pattern across the landscape. It generally represents small areas that might include household agriculture and traditional cattle ranching (Almeida et al., 2016). | Mosaic of uses |
| Noncombustible areas | Areas with low vegetation density (fuel accumulation) or high humidity and water levels. This class was excluded from the analysis. | Mangrove; Hypersaline Tidal Flat; Wetland; Rocky Outcrop; Non vegetated area; Water |

### 2.2 Fire data

To obtain maps of fires in the ecoregions of the AF, we used the MapBiomas platform – Fire Collection 2.0, which includes annual records of fire scars from 1985 to 2022 (MapBiomas, 2024). Similar to the LULC maps, this project also identifies burned areas through analyses of Landsat images with 30-m pixels, using the Normalized Burn Ratio (NBR) index (see more details in https://brasil.mapbiomas.org/metodo-mapbiomas-fogo/). To ensure greater accuracy in identifying the parts of the LULC that burned, we aligned our fire maps with our LULC data collection, which spans from 1986 to 2021 with a five-year interval, selecting fire maps from years that corresponded to one year after each LULC map. Consequently, our fire maps, also selected at five-year intervals, span from 1987 to 2022. The minimum fire extent was five hectares (see Pereira and Santos, 2003; Moreira et al., 2009; Silva et al., 2009; Carmo et al., 2011), with each fire assigned to an ecoregion of the AF.

### 2.3 Data analysis

To determine the relative fire-proneness of each LULC type, we employed a selection ratio-based approach. This method involved comparing, for each fire, the proportion of each LULC type within the burned area to its proportion in the total area (burned and unburned) available for burning. In our study, a landscape unit is therefore defined as the spatial mosaic of different LULC types within the total area available for burning.

To define the total area available for burning, several methods have been proposed in the literature. For example, Moreira et al. (2009) and Silva et al. (2009) defined circular buffers centered on each fire, with a radius corresponding to the size of the largest fire in the respective ecoregion, which was expected to approximate fire shapes in the absence of external influences. Alternatively, Oliveira et al. (2014) tailored buffers to the specific characteristics of each fire’s perimeter, using a shape that mirrored the fire but was approximately twice its size. Our approach aligns with Oliveira et al. (2014) in creating buffers that mirror the shape of each fire, but we opted for a buffer size of approximately four times the fire area based on preliminary analyses indicating that a larger buffer was necessary in our study region to capture variations in LULC while minimizing overlap between fires. This method was originally proposed for the study of resource selection by animals (Manly et al., 1993) and has been applied in fire research studies (e.g., Moreira et al., 2009; Silva et al., 2009; Carmo et al., 2011; Oliveira et al., 2014).

The selection ratio (*w_i_*) for a given LULC type *i* is an index estimated as *w*_i_ = *o*_i_/π_i_ (Manly et al., 1993), where *o_i_* is the proportion of a burned patch belonging to LULC type *i* (estimated from the area consumed by fire) and π_i_ is the proportion of available area belonging to LULC type *i* (estimated from both the burned patch and the surrounding buffer). Selection ratios range from 0, if the LULC type is available but not consumed by the fire (*o_i_* = 0), to a large value (in theory ∞), in the situation where the fire consumes the only tiny patch of a given LULC type that was available to burn (π_i_ approaches 0) (Moreira et al., 2009). To avoid these extreme *w* values in very small areas, we only analyzed LULC patches larger than 1 ha. If a given LULC type is burned in proportion to its availability, then *w*=1. If *w*>1, the LULC type is considered more fire-prone than expected by chance. Conversely, if *w*<1, the LULC type is considered less fire-prone than expected by chance. Since we were interested in the fire-proneness of LULC types per ecoregion, we omitted fires that, with their buffers, overlapped more than one region. A total of 40,128 burned patches were considered for analysis.

Next, we averaged values for each LULC type across the fires within a given ecoregion and estimated 95% confidence intervals. We considered differences between selection ratios for different classes significant when their respective confidence intervals did not overlap. Additionally, if the confidence interval of *w* did not include 1, we considered the class significantly more or less prone to fire than expected by chance.

## 3. RESULTS

Of the 40,128 fires detected and analyzed between 1987 and 2022, 44% occurred in the SSF ecoregion, 21% in CA, and the lowest percentages were in MOF (9%) and DSF (7%) (Table 2). The mean fire size was 23 ha, ranging from 15 ha in MOF to 27 ha in SSF. Mosaic of uses, secondary forests, and old-growth forests, in that order, were the LULC types with the highest available areas across regions. The region with the highest proportion of old-growth forests coincides with the area with the most fires, the SSF. CA had the highest proportion of secondary forests.

**Table 2.** Number of fires larger than 5 hectares and mean fire size (ha) by ecoregion and the percentage of area occupied by each type of LULC.

| Ecoregion | No. of fires | Mean fire size (ha) | LULC types |  |  |  |  |  |  |  |
| --- | --- | --- | --- | --- | --- | --- | --- | --- | --- | --- |
|  |  |  | Old-growth forests | Openlands | Temporary crops | Perennial crops | Plantation forests | Mosaic of uses | Secondary forests | Noncombustible areas |
| Semideciduous seasonal forest | 17587 | 20.9 | 27.3 | 11 | 2.3 | 1.4 | 0.6 | 33.3 | 19.3 | 4.9 |
| Deciduous seasonal forest | 2923 | 20.7 | 18.8 | 25.4 | 0.1 | 0.5 | 0.9 | 26.9 | 26.3 | 1.2 |
| Dense ombrophilous forest | 7777 | 16.3 | 22.8 | 12 | 2.4 | 0.6 | 1.5 | 33.4 | 21.2 | 6.2 |
| Mixed ombrophilous forest | 3472 | 10.6 | 22.8 | 10 | 5.7 | 0.6 | 3.4 | 29 | 24.4 | 4.1 |
| Contact areas | 8369 | 16.8 | 15.9 | 13.8 | 3.5 | 1.5 | 1 | 21.1 | 38.5 | 4.7 |

### 3.1 Fire-proneness across ecoregions

Across the ecoregions of the Brazilian AF, secondary forests were the most fire-prone type of LULC, while old-growth forests were the least fire-prone (Fig. 2). Secondary forests burned 61% more than expected by chance (*w* = 1.61), whereas old-growth forests burned 57% less (*w* = 0.43). Besides secondary forests, only the mosaic of uses LULC type was significantly prone to fire, with a selection ratio 5% higher than expected. Like old-growth forests, the LULC types openlands, temporary crops, and planted forests were also significantly less prone to fire than expected, with selection ratios between 31% and 35% below expected. Perennial crops were the only type of LULC to burn according to availability across AF’s ecoregions (confidence interval of *w* included 1).

**Figure 2.**
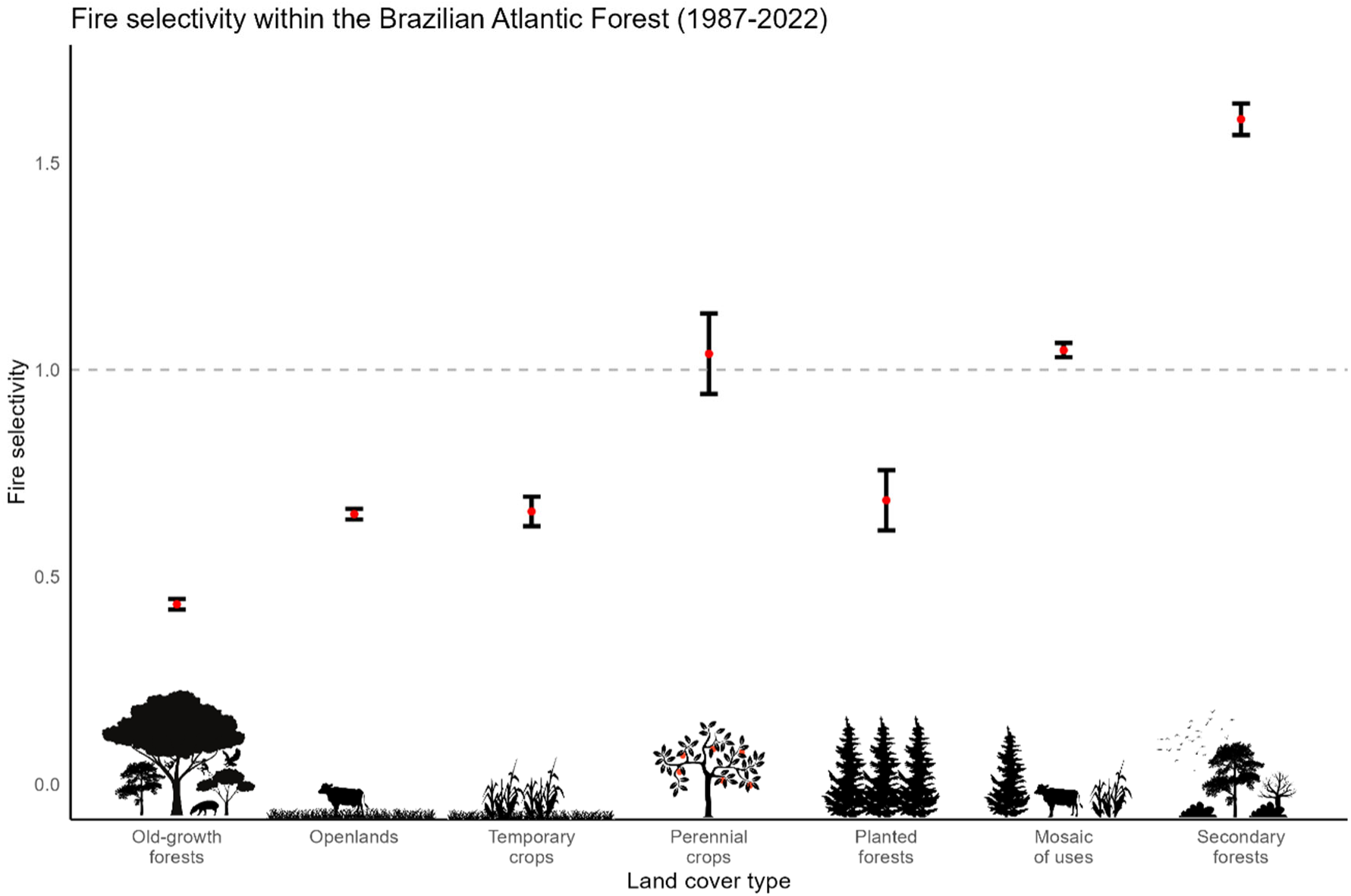
Mean selection ratios (*w*) with 95% confidence intervals for LULC types across ecoregions of the Brazilian Atlantic Forest. *w* = 1 indicates the LULC type burns according to availability; *w* < 1 means it burns significantly less, and *w* > 1 means it burns significantly more than expected by chance.

### 3.2 Fire-proneness by ecoregion

As hypothesized, old-growth forests had low fire proneness across ecoregions (Fig. 3), burning between 33% and 66% less than expected by chance, depending on the region. Except for the DSF region, this LULC type had the lowest selection ratio in all other regions. In contrast, with selection ratios between 46% and 87% above expected, secondary forests tended to have the highest fire proneness across regions, except for the DOF. Openlands had low fire-proneness, burning 20% to 59% less than expected, except in MOF where they burned according to availability. As expected, other human-impacted LULC types showed variable fire proneness, depending on the ecoregion.

**Figure 3.**
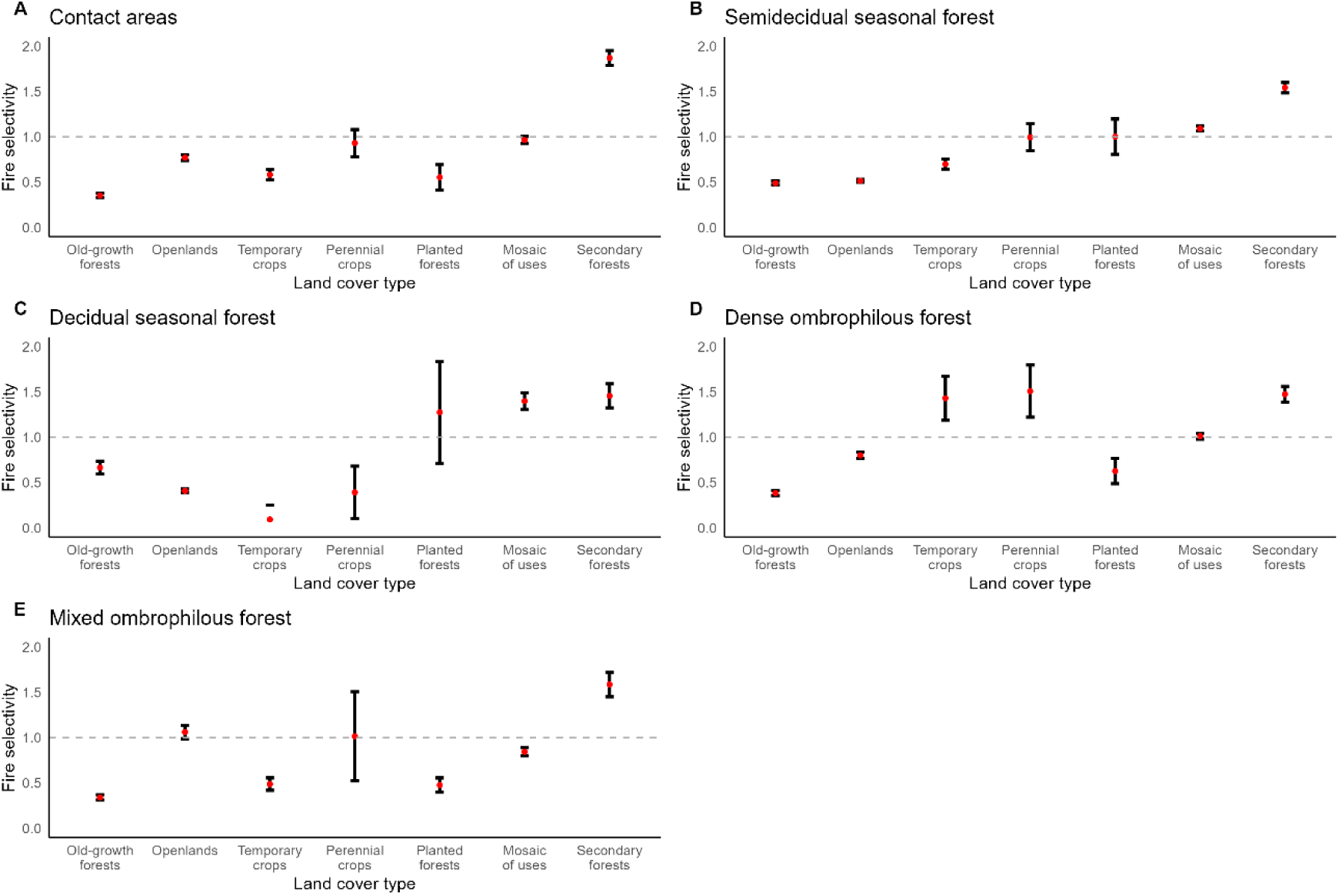
Mean selection ratios (*w*) with 95% confidence intervals for LULC types in each ecoregion of the Brazilian Atlantic Forest. Refer to Fig. 2 for more explanations.

### 3.3 Fire-proneness over time

The fire-proneness of old-growth forests remained consistently low over the 35-year study period (Fig. 4), showing no pattern of increase or decrease. Likewise, openlands and temporary crops generally had low fire-proneness, with no clear change over time, although temporary crops burned according to availability in 1992. Perennial crops, which generally burned in proportion to availability until 2012, became less prone to burning than expected from that year onwards. Forest plantations consistently had low fire-proneness, except in 1992 and 2002 when their fire-proneness was neutral. Mosaic of uses showed mixed fire-proneness over the years but shifted to a significantly lower than expected fire-proneness from 2012 onwards. It burned in proportion to its availability in 1992 and 2012, had high fire-proneness in 1987 and 1997–2007, but had low fire-proneness in 2017 and 2022. Secondary forests followed the hypothesized trend of decreasing fire-proneness, being significantly prone to fire until 2012, then burning in proportion to their availability from that year onwards.

**Figure 4.**
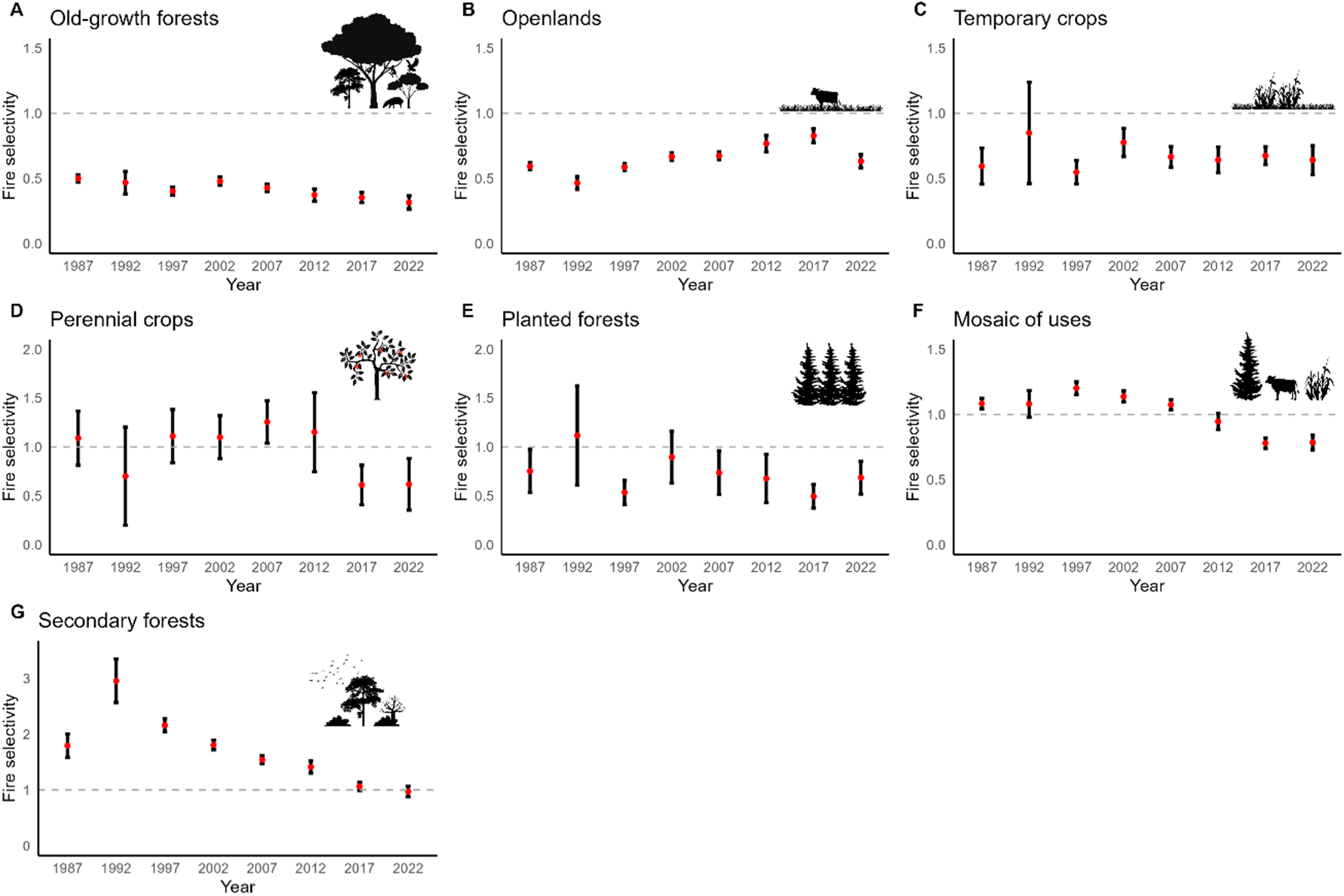
Mean selection ratios (*w*) with 95% confidence intervals for LULC types across ecoregion of the Brazilian Atlantic Forest from 1987 to 2022. Refer to Fig. 2 for more explanations.

## 4. DISCUSSION

Our analysis spanning over 35 years of fire-LULC interactions in the Brazilian AF ecoregions provides insight into the relative fire-proneness of different LULCs in a tropical biome. This is critical as fire becomes a growing issue both worldwide and in these biomes. Across all ecoregions, our results confirmed the low fire-proneness of old-growth tropical forests, in contrast with the high fire-proneness of secondary forests.

Other LULCs showed more variable fire-proneness, with some differences between ecoregions. Furthermore, fire-proneness has changed over time in several LULCs, suggesting that these trends may be driven by natural ecological succession, climatic conditions, or human actions potentially influenced by changes in Brazilian environmental legislation. Notably, we found a decline in the fire-proneness of secondary forests over time, likely due to their lower susceptibility as the average age of these patches in the landscape increases, resulting in characteristics increasingly similar to old-growth forests.

### 4.1 Fire-proneness in the Brazilian Atlantic Forest

By confirming the low fire-proneness of old-growth tropical forests, our results are consistent with studies indicating a reduced risk and density of fires in undisturbed forests within the AF (de Assis Barros et al., 2022; Guedes et al., 2020; Singh & Huang, 2022). Despite the significant fragmentation and deforestation in the AF (Ribeiro et al., 2009; Vancine et al., 2024), we demonstrated that, in general, old-growth forest fragments still remain the LULC type least likely to burn compared to all others. The low fire-proneness of tropical old-growth forests can be attributed to their retention of transpired moisture and the trees’ substantial contribution to forest humidity, which limits fire ignition and spread (Cochrane, 2003a; Cochrane et al., 1999). Interestingly, our data showed that the likelihood of old-growth forests burning less than expected across regions was similar to that of secondary forests burning more than expected, both at approximately 60%.

The high fire-proneness of secondary forests was anticipated, as they exhibit high fuel load, low moisture content, and high radiation and temperature in the understory (Gutiérrez-Vélez et al., 2014; Hasselquist et al., 2010; Kyereh et al., 2007). Additionally, the spatial arrangement of these areas likely influences the fire spread, as they may be closer to anthropized areas and sources of ignition. This result is also consistent with previous works in the Amazon Forest that found higher fire activity in degraded forests (Balch et al., 2015; Brando et al., 2014, 2016; Cochrane et al., 1999). Research further suggests that, because secondary forests are often viewed as having low economic value, farmers may be less motivated to prevent them from burning when nearby fires pose a threat (Sorrensen, 2000). Other studies also show a link between deforestation fires to expand agricultural areas and the high fire-proneness of secondary forests (Pivello et al., 2021). Despite Brazilian legislation prohibiting fires, except in specific situations, farmers still burn secondary forests in pastures and agricultural areas to prevent forest encroachment. Furthermore, these areas are often re-cleared within the first few years following forest establishment (Piffer et al., 2022).

Among the LULC types significantly less prone to fire, the results for openlands, where pastures are included, were noteworthy. Despite pastures having the largest burned areas in Brazil (Alencar et al., 2022; Moreira de Araújo et al., 2012) and being generally associated with fire risk (Guedes et al., 2020; Gutiérrez-Vélez et al., 2014; Herrmann et al., 2023), our findings indicate lower-than-expected fire proneness. This shows the advantage of our selection ratio-based approach, as this method allows a distinction to be made between when a LULC type burns extensively due to its abundance in the landscape and when it burns disproportionately more than its availability. On the contrary, previous works have generally highlighted the large representativeness of pastures within the burned areas, overlooking the total available area that could have potentially burned in the landscape, which is usually also substantial. Indeed, although pastures occupy around 30% of the AF (Dos Santos et al., 2022; Souza et al., 2020), our results show that their fire proneness is likely lower than expected when availability per fire is taken into account. The high association of fire with pastures found in other studies may not be due to a positive selectivity compared to other land covers, but rather to the frequent illegal use of fire for cleaning and renewal pastures (Moreira de Araújo et al., 2012). It should be noted that although our results indicate lower fire proneness for pastures than expected by chance, pasture fires still pose a threat to the AF by contributing to the spread of fires to other fire-prone LULCs (Guedes et al., 2020).

Management-related aspects of the other LULC types that were significantly less prone to fire, such as temporary crops and planted forests, or that burned according to availability, like perennial crops, should be stressed. Managed croplands usually exhibit low fuel load and high moisture content due to irrigation (Duguy et al., 2007). Planted forests, on the other hand, are often intensively managed by private companies to avoid fire hazards (Guedes et al., 2020; Mirra et al., 2017). Additionally, these LULCs are typically closer to urban areas, roads, and scattered populations. This proximity may result in more frequent fire occurrences (dos Santos et al., 2019; Barros et al., 2022), but it also facilitates faster fire detection and easier firefighting efforts, reducing burned extent and fire proneness (Moreira et al., 2009).

Lastly, it was to be expected that mosaic of uses would be prone to fire across ecoregions. These areas typically consist of small properties where slash-and-burn practices are employed to renew the land and eliminate waste. Subsistence farmers and indigenous populations often use these fires, confining them to small areas with localized impacts (Pivello et al., 2021). However, there is a significant risk that these fires could accidentally spread to nearby fire-prone LULCs. Under climate change scenarios, the risk of fire increases, especially when traditional agricultural fire practices result in accidents or loss of control (Pivello et al., 2021). It is crucial to develop strategies that balance traditional practices with modern fire management to mitigate these risks effectively.

### 4.2 Fire-proneness by ecoregion

Although it was hypothesized and therefore unsurprising that the old-growth forests of the AF exhibit consistently low fire-proneness with minimal variation between ecoregions, it is nonetheless remarkable that this low fire-proneness remains uniform across regions characterized by dry seasons, such as SSF and DSF, as well as in regions with evergreen vegetation, like ODF. These results reinforce the reduced fire-proneness of tropical forests (Cochrane, 2003; Cochrane et al., 1999), even under disparate regional conditions. Yet, previous studies have also shown that altitude, mean annual temperature, and drought severity can be positively associated with fire occurrence and lead to variations in the area burned in the AF (Abreu et al., 2022; de Assis Barros et al., 2022; de Santana et al., 2020; Herrmann et al., 2023; Singh & Huang, 2022).

In the AF, regions with varying economic development can show different success rates in restoration programs (Piffer et al., 2022), affecting fire prevention and control. In the Brazilian Amazon, studies show that federal and state agencies, with distinct institutional capacities, implement diverse fire management practices, contributing to regional differences in fire management (Fonseca-Morello et al., 2017). In Portugal, research also shows that regions with different climatic and environmental conditions present LULCs with different fire-proneness (Moreira et al., 2009). In our study, this is likely why the LULC openlands type, which generally had low fire-proneness in most AF ecoregions, exhibited higher fire-proneness, burning according to its availability, in the MOF region. This result is likely influenced by both the environmental and cultural context of the MOF ecoregion. This ecoregion is characterized by higher average altitudes and lower temperatures (IBGE, 1992), and is primarily concentrated in southern Brazil. Unlike the southeastern and northeastern regions, where intensive agriculture predominates, southern Brazil has a larger number of properties engaged in family farming. In these settings, the use of fire in pastures is a traditional practice (Carvalho & Andrade-Filho, 2019). Additionally, some southern states, such as Rio Grande do Sul, grant municipalities the authority to authorize and supervise the use of fire in pastures (Law No. 13,931, 2012). These regional differences are likely related to the smaller average fire sizes observed in this ecoregion compared to others.

Despite the environmental, cultural, social, and economic differences between ecoregions, the secondary forest LULC type burned more than expected by chance in all AF regions, as predicted. Not even the consistently high rainfall in evergreen ecoregions such as ODF (Oliveira et al., 2000) reduces the fire-proneness of secondary forests, highlighting the vulnerability of this LULC to such disturbance. Still, our results show that secondary forests are almost twice as fire-prone in CA as in DSF, with selection ratios 87% and 46% higher than expected, respectively. This may be due to the fact that CAs are mainly transition zones between the AF biome and the Brazilian Cerrado. Unlike the AF, the Brazilian Cerrado is a fire-dependent system, with ecological communities adapted to this disturbance (Pivello et al., 2021). As such, the greater proximity of CA’s secondary forests to the Cerrado is likely to explain their greater fire proneness.

Likewise, as in the Cerrado, where cattle ranching is the leading cause of fire (Pivello et al., 2011), secondary forests in the CA regions are also burned to expand pasturelands. Worryingly, fires in secondary forests can promote the growth of herbaceous species and lead to the replacement of forests by non-woody vegetation types. This shift is concerning, as positive feedback loops with fire can maintain these new vegetation types, which is a growing issue in tropical forests worldwide (Pivello & Coutinho, 1996; Sansevero et al., 2020).

### 4.3 Fire-proneness of the Brazilian Atlantic Forest over time

Our results also confirmed the hypothesis that AF old-growth forests would exhibit consistently low fire-proneness over time, with minimal variation across the sampling years. Not even the inclusion of a major El Niño year (1996–1997) among the years sampled altered this consistency, despite major El Niño years being associated with extreme droughts and a greater potential for fires to occur and spread (Aragão et al., 2018; Marengo et al., 2018). In fact, this El Niño event does not seem to have altered the fire-proneness in 1997 or the chronosequential fire-proneness pattern for any LULC type studied. However, the years sampled did not include other extreme climatic events, such as the major La Niña (characterized by high precipitation) in 1988–1989 or the major El Niño in 2015-2016. Therefore, it remains possible that some LULC types had unusual fire-proneness patterns during these events. Future studies on fire-landscape interactions in the AF should include specific climatic drivers, such as variations in precipitation, temperature anomalies, and the occurrence of extreme weather events associated with major El Niño and La Niña phenomena.

The hypothesized decreasing fire-proneness trend of secondary forests over time was confirmed and the result can be discussed in light of two main reasons. First, the early data from the MapBiomas collection that we used to obtain the LULC maps only included young secondary forests. Over time, the collection encompassed a broader range of ages for this LULC type, leading to an increase in the average age of secondary forest patches included in the annual maps. Thus, the reduced fire-proneness of secondary forests in more recent years, burning in proportion to availability from 2012 onwards, is likely partly due to the increased age of these forests. Forest succession leads to significant environmental changes, with early stages having higher temperatures and lower humidity in the understory, transitioning to cooler and more humid conditions in later stages (Ray et al., 2010; Lebrija-Trejos et al., 2011). However, the approximate maximum age of the secondary forest patches in our data in 2022, 35 years, combined with the existence of younger patches, means that this LULC type has not yet reached the tipping point of having characteristics close enough to old-growth forests to become equally less prone to fire. Future studies must analyze the age of each patch of secondary forest so that we can understand more precisely after how long this type of LULC becomes less prone to fire than expected by chance.

The second reason for the decreasing fire-proneness is that early-stage secondary forests are often burned to prevent encroachment on agricultural areas. In the AF, regenerated forests in pastures and areas of shifting cultivation—where land is alternately farmed and left to regenerate—are frequently re-cleared before reaching advanced succession stages (Piffer et al., 2022), thus avoiding stricter protection under the Atlantic Forest Law (Law No. 11.428/2006). This law highly restricts cutting native vegetation, allowing it only in initial and medium regeneration stages with appropriate authorization, provided it follows sustainable management criteria. Cutting vegetation over 5 years old is prohibited. On the other hand, legislation changes may also be behind the shift to low fire-proneness from 2012 onwards for the perennial crops and mosaic of uses LULC types, after the approval of the New Brazilian Forest Code (Law n° 12.651/2012).

Firefighting may well have become more effective in these LULCs after the enactment of this law, which devotes a chapter to the prohibition and control of wildfires.

### 4.4 Management implications

Our results show that secondary forests have disproportionately burned the most across all regions of the AF in the past few decades, emphasizing the need for fire management strategies to carefully address this LULC. This is even more crucial considering that numerous restoration programs have restored millions of hectares of native vegetation, primarily as secondary forests, in recent decades (dos Santos et al., 2019; Brancalion et al., 2019; Vancine et al., 2024). It is clear that a trade-off must be established between the benefits of increasing secondary forest areas and the costs of protecting them, especially in the early years, to avoid a greater risk of fire in the landscape. Fire will lead to the interruption of forest succession to mature forests (Cochrane, 2001; Cochrane et al., 1999; Sansevero et al., 2020). Implementing effective fire prevention and management measures in secondary forests will be necessary to safeguard the restoration efforts and ensure their long-term success (dos Santos et al., 2019). Some authors simply argue that in the face of deforestation fires, which are illegal but still used in secondary forests, law enforcement should be the primary response, defending that fines should then be used to fund law enforcement (Pivello et al., 2021).

By showing such a large discrepancy in fire-proneness between secondary forests and other LULC types, our data also suggest incorporating integrated landscape approaches from fire-prone landscapes in Mediterranean regions and North America (Pivello et al., 2021). One such approach is ‘fire-smart management’, which involves managing the entire landscape to reduce fire risk. In this context, implementing low fire risk matrices composed of LULC types that our results have shown to be less prone to fire around secondary forests can be an effective strategy to prevent the spread of fire. For example, there is evidence that eucalyptus plantations provide some protection against fire, because plantation owners seek to suppress and control fires that could otherwise harm their eucalyptus investments (Guedes et al., 2020). Using this type of LULC around more fire-prone patches, including mosaic of uses, may be an option to consider.

The fire-proneness of secondary forests in the AF also highlights the need to increase their perceived value by local communities to prevent them from burning (Guedes et al., 2020). This could involve enriching younger forest patches (e.g., up to 5 years old) with fruit or timber trees and offering payments for ecosystem services such as carbon sequestration, water regulation, and habitat restoration (Guedes et al., 2020). Some authors suggest that financial institutions could offer better loan conditions and payment rates to farmers with a good fire governance record (Nepstad et al., 2014). Similarly, commodity buyers may provide favorable terms to farmers that adhere to the rules (Pivello et al., 2021). Lastly, the AF should contain programs dedicated to forecasting and monitoring wildfires, like those in the Amazon Forest. In this scenario, plans to combat and prevent fires should be developed well in advance, taking into account alerts for areas with high deforestation rates, especially those with secondary forests (Pivello et al., 2021).

## 5. Conclusion

This study used a selection ratio-based approach in 40,128 fires spanning over 35 years to evaluate the relative fire-proneness of LULC types in a tropical biome for the first time. This method compares, per fire, the proportion of each burned land cover type with its availability nearby. Our findings quantified the variation in fire-proneness across LULC types within the Brazilian AF, highlighting that the likelihood of secondary forests burning more than expected was approximately 60%, mirroring the 60% likelihood of old-growth forests burning less than expected. Interestingly, pastures, included in openlands, exhibited lower-than-expected fire proneness despite their large burned areas in Brazil. Secondary forests burned more than expected in all AF regions, with a particularly high fire-proneness in CA, transition zones between the AF and the Cerrado, a fire-dependent system. The observed trend of decreasing fire-proneness in secondary forests over time likely reflects both the increasing age and maturity of these forests and changes in land management practices.

We emphasize the necessity for tailored fire management strategies that address the unique vulnerabilities of secondary forests, particularly in the context of ongoing restoration efforts aimed at increasing forest cover. Effective measures, including the implementation of ‘fire-smart management’ practices and enhancing the perceived value of secondary forests among local communities, are crucial for mitigating fire risks. Integrating these strategies with robust law enforcement and incentive-based approaches can bolster fire prevention and control, ensuring the long-term success of restoration programs. Our analysis provides a framework for understanding fire-landscape dynamics in tropical forests and offers actionable insights for policymakers, conservationists, and land managers working to safeguard these biomes from the escalating threat of wildfires.

## Acknowledgments

This study was financed in part by the Coordenação de Aperfeiçoamento de Pessoal de Nível Superior - Brasil (CAPES) - Finance Code 001 (Grant 88887.889095/2023-00). BA (Grant 140361/2021-9) and AJP (Grant 316032/2023-9) were funded by CNPq—Conselho Nacional de Desenvolvimento Científico e Tecnológico. PGV (CE3C DOI 10.54499/UIDB/00329/2020; CHANGE DOI 10.54499/LA/P/0121/2020) was funded by FCT—Portuguese Science and Technology Foundation.

## Authors’ contributions

**Bruno Adorno:** Conceptualization, Investigation, Formal analysis, Writing - Original Draft, Writing – Editing. **Augusto Piratelli:** Writing - Review & Editing, Supervision. **Erica Hasui:** Writing - Review & Editing. **Milton Ribeiro:** Writing - Review & Editing. **Pedro Vaz:** Conceptualization, Formal analysis, Writing - Original Draft, Writing - Review & Editing, Supervision.

## Notes

### Competing Interest Statement

The authors have declared no competing interest.

### Summary of Updates

Revised version incorporating feedback from two anonymous reviewers after the first round of revisions.

